# BRD4 phosphorylation regulates the structure of chromatin nanodomains

**DOI:** 10.1101/2024.09.03.611057

**Authors:** Clayton Seitz, Donghong Fu, Mengyuan Liu, Hailan Ma, Jing Liu

**Affiliations:** Department of Physics, Indiana University, Indianapolis, IN 46202, USA; Department of Physics and Astronomy, Purdue University, West Lafayette, IN 46202, USA; Melvin and Bren Simon Comprehensive Cancer Center, Indiana University, Indianapolis, IN 46202, USA

## Abstract

The interplay between chromatin structure and phase-separating proteins is an emerging topic in cell biology with implications for understanding disease states. Here, we investigate the functional relationship between bromodomain protein 4 (BRD4) and chromatin architecture. By combining molecular dynamics simulations with live-cell imaging, we demonstrate that BRD4, when mutated at specific N-terminus sites, significantly impacts nucleosome nanodomain (NN) organization and dynamics. Our findings reveal that enhanced chromatin binding activity of BRD4 condenses NNs, while both loss or gain of BRD4 chromatin binding reduced diffusion of single nucleosomes, suggesting a role for BRD4 in the regulation of nanoscale chromatin architecture and the chromatin microenvironment. These observations shed light on the nuanced regulation of chromatin structure by BRD4, offering insights into its role in maintaining the nuclear architecture and transcriptional activity.

## INTRODUCTION

The cell nucleus is a densely packed environment with chromatin comprising a dominant component. Emerging research supported by advanced imaging and sequencing methods have revealed that chromatin is dynamic and highly compartmentalized^1,2^. The compartmentalization of chromatin with other intranuclear components is therefore an efficient strategy to ensure precise spatial and temporal coordination of complex dynamics towards the regulation of intracellular activities and cellular functions. On one hand, nanoscale chromatin movement (chromatin motion) modulates the interaction of DNA with regulatory molecules (chromatin accessibility), thereby influencing global gene expression patterns^40-45^. A growing number of phase separated nuclear bodies including transcriptional condensates^3–9^, nuclear speckles^10–14^, and DNA damage repair foci^15–20^ have been identified. However, the interplay of phase separated condensates with the underlying chromatin structure remains poorly understood.

Transcriptional condensates are a suitable model to study the kinetic and thermodynamic contributions of chromatin substrate binding, as the ability of transcriptional activators to both condense and bind chromatin is well established^3,21–24^. For example, BRD4 protein is a well-studied transcriptional activator that undergoes phase separation^25^, localizes to acetylated chromatin sites^26^, recruits pTEF-b^27^, and initiates transcription of key genes involved in signal response, immunity, and oncogenesis^27^. The BRD4 long isoform is characterized by structured N-terminal tandem acetyl-lysine binding bromodomains and an extra-terminal domain, connected by intrinsically disordered regions^28^

Recent evidence suggests a phosphorylation-dependent binding mechanism of BRD4 to acetylated chromatin^29,30^. However, the binding kinetics of BRD4 proteins with chromatin cannot be directly transferred to studying the interplay of transcriptional condensates with the underlying chromatin structure, and the mechanistic role of chromatin dynamics in transcriptional regulation remains unclear. Therefore, we speculated that BRD4 phosphorylation can modulate a multivalent binding interaction between the transcriptional condensate with NNs, making BRD4 phosphorylation necessary for the maintenance of NN structure and dynamics.

## RESULTS

### Colocalization of BRD4 mutants with nucleosome nanodomains

To address the role of BRD4 binding and phase separation on chromatin structure, we express FLAG-tagged BRD4 mutants in HeLa cells and measure their effects on chromatin organization. Perhaps the most fundamental of BRD4 functions is the ability to bind to acetylated chromatin through bromodomain 1 (BD1) and bromodomain 2 (BD2) that are in tandem within the N-terminal part of the protein. BRD4 inhibitors such as (+)-JQ1 competitively bind to the acetyl-binding pocket of BRD4, displacing BRD4 from chromatin^28^. It is also well known that BRD4 association with acetylated chromatin is enhanced by casein kinase II (CK2)-mediated phosphorylation of seven N-terminus phosphorylation sites (NPS), followed by intramolecular rearrangement^32^ of BRD4 protein and/or BRD4 dimerization^32,33^. Therefore, we express a constitutively phosphorylated BRD4 mutant with 7 aspartate substitutions in the NPS region (7D mutant), a constitutively unphosphorylated mutant with 7 alanine substitutions in the NPS region (7A mutant), and bromodomain-deactivated (BD mutant) (Figure 1a). We find similar bulk expression levels of all mutants (Figure 1b). Colocalization analysis of BRD4 mutants with NNs using the nearest neighbor distance distribution showed an obvious colocalization of these mutants with NNs. The 7D mutant showed the closest proximity to NNs relative to wild type, followed by BD and 7A mutants (Figure 1d). This result is consistent with known dependence of BRD4 chromatin binding on phosphorylation state.

**Figure 1:**
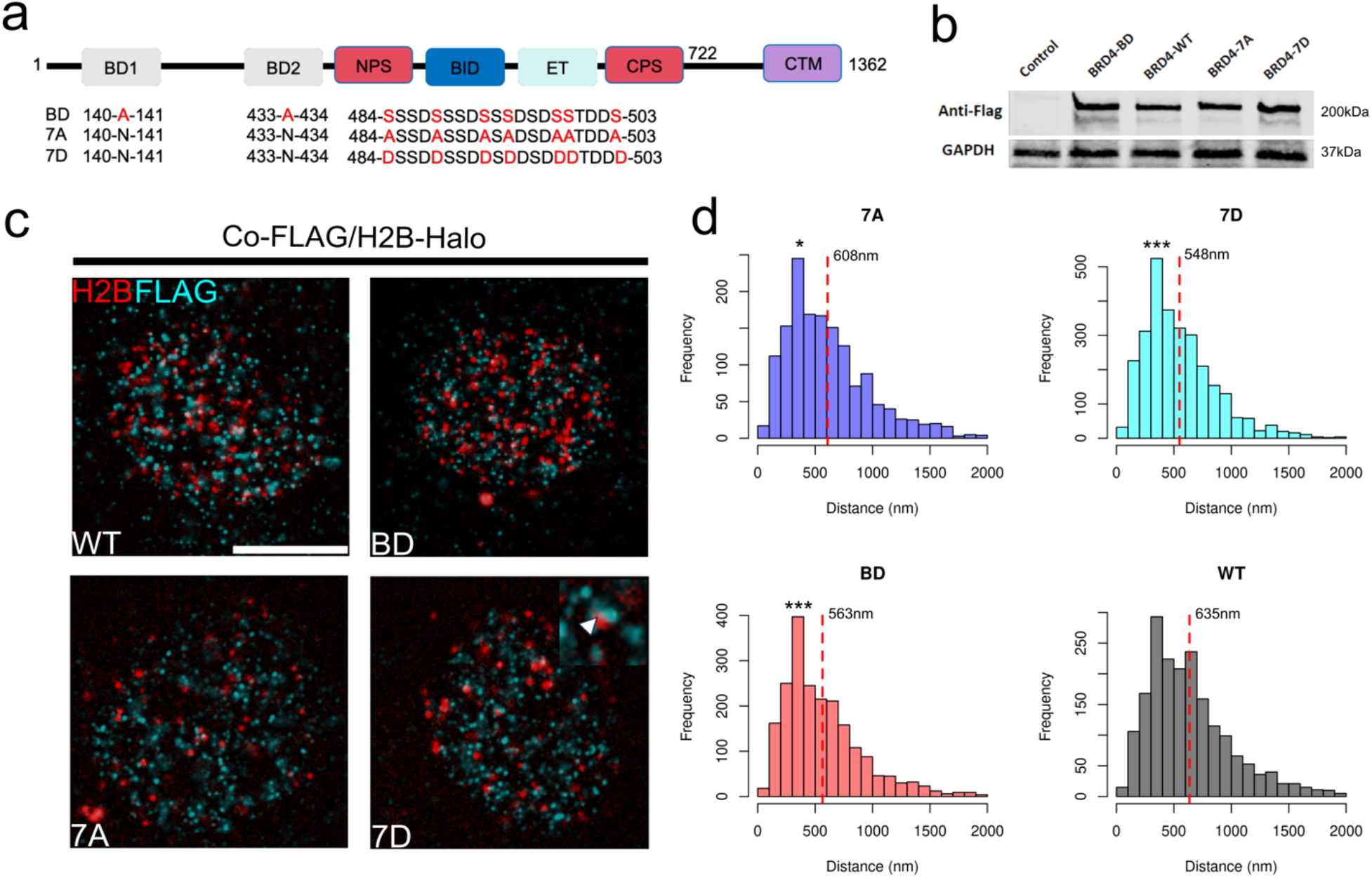
BRD4 mutants colocalize with nucleosome nanodomains. (a) Schematic of the BRD4 protein sequence and mutations shown in red. (b) Anti-FLAG western blot of bulk expression of BRD4 mutants. (c) Combined immunofluorescence of FLAG-tagged BRD4 mutants with JF646-tagged H2B-Halo. (d) Histograms of nearest neighbor distances of H2B puncta to FLAG puncta over N=10 cells for each mutant. Red dashed lines are drawn at the mean, statistical significance determined by Welch t-test for each mutant relative to wild-type (WT) * p<0.05, *** p<0.001

### Phase-separated BRD4 condensates regulate chromatin structure and dynamics

To assess the functional role of BRD4 in maintaining the nucleosome environment, we interrogated the dynamics of NNs, as well as their structure, in the presence of BRD4 mutants in live Hela cells. Histone H2B was tagged with HaloTag^34^ (H2B-Halo), to which a fluorescent ligand JaneliaFluor646 (JF646) can bind specifically in a living cell. Low concentrations of JF646 were used to obtain sparse labeling of nucleosomes for single-nucleosome imaging (Figure 2a,b). JF646-labeled nucleosomes in HeLa cells were recorded at 10fps (∼200 frames, 20 s total) and a reduced diffusion coefficient was measured in cells expressing 7A, 7D, and BD mutants, with respect to cells expressing the wild-type protein (Figure 2c). Meanwhile, HeLa cells exposed to (+)-JQ1 in DMSO for 8h showed an increase in nucleosome dynamics with respect to DMSO alone (Figure 2d). We then conducted super resolution imaging of NNs using direct stochastic optical reconstruction microscopy (dSTORM) by promoting JF646 fluorescence intermittency with a cysteamine buffer (Figure 3a,b). JF646 is known to exhibit a transient fluorescent state lasting tens to hundreds of milliseconds and stable dark state lasting hundreds of milliseconds to seconds. Two color imaging of H2B-Halo-JF646 and GFP-tagged BRD4 shows that BRD4 and NNs form complementary biomolecular condensates in the nucleus, consistent with current models of a BRD4 chromatin reading mechanism (Figure 3c). Ensemble averages of Besag’s L-function showed an increase in nucleosome clustering in cells expressing the 7D BRD4 mutant, while all other groups were consistently indistinguishable from WT cells (Figure 3e). HeLa cells exposed to (+)-JQ1 in DMSO for 8h showed a reduction in nucleosome clustering with respect to DMSO alone (Figure 3f).

**Figure 2:**
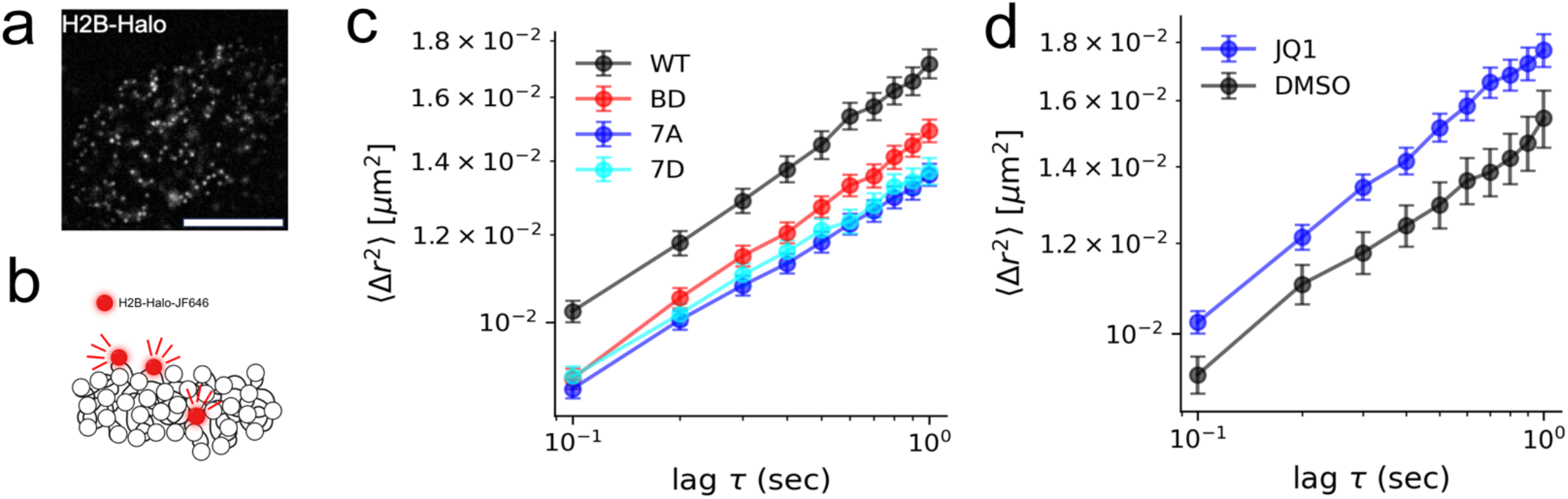
BRD4 mutants reduce single nucleosome dynamics in living cells. (a,b) Sparse labeling of single nucleosomes in a living HeLa cell (scalebar 5um). (c) Average mean squared displacement (MSD) of single nucleosomes for wild-type and mutated BRD4, error bars represent the standard error of the mean. (d) Average MSD of single nucleosomes after 8h exposure to 1uM (+)-JQ1 in DMSO and DMSO only, error bars represent the standard error of the mean

**Figure 3:**
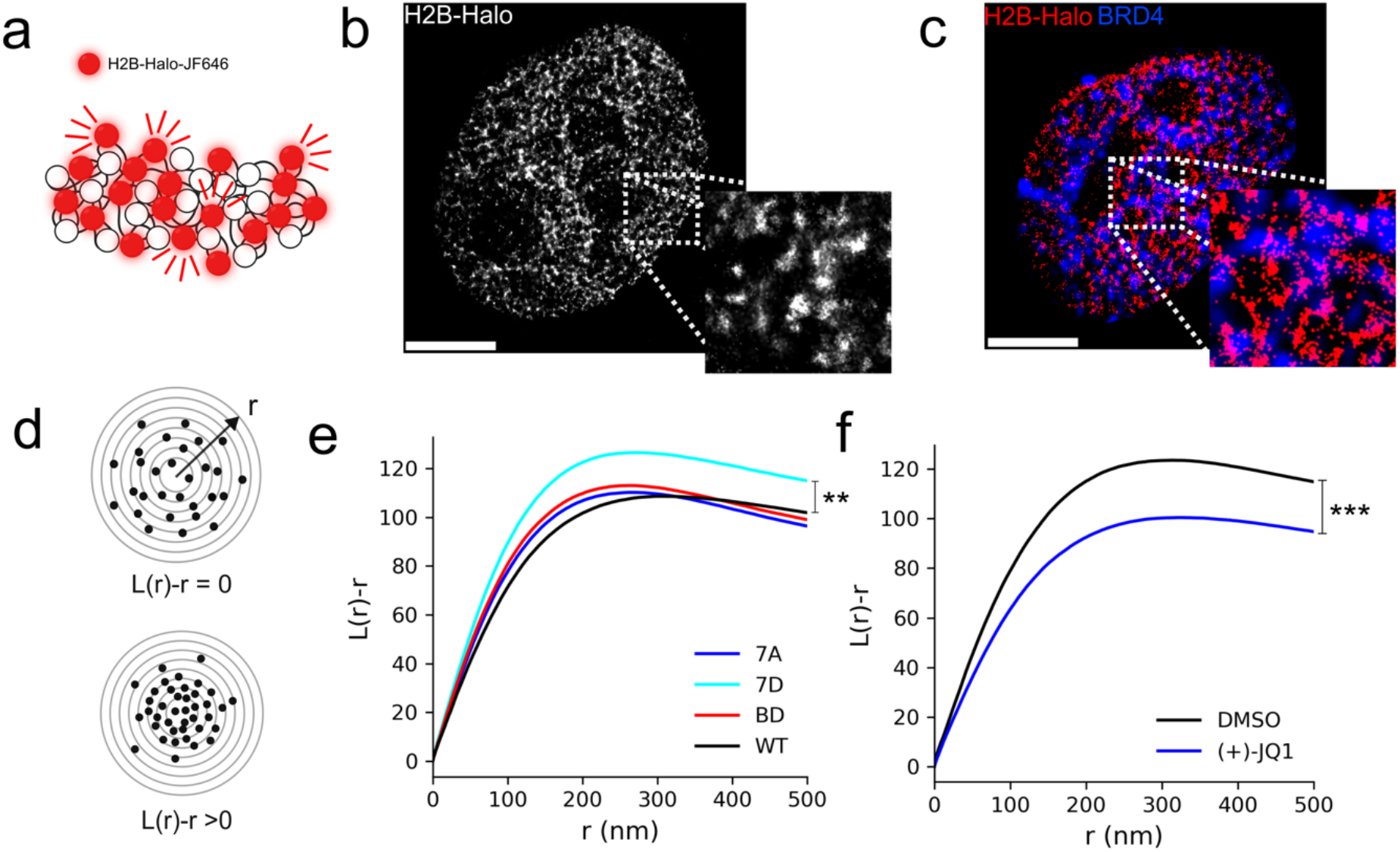
Expression of constitutively phosphorylated BRD4 compacts nucleosome nanodomains. (a,b) Direct stochastic optical reconstruction microscopy (dSTORM) imaging strategy of single nucleosomes and an example super-resolution image. (c) Two-color image of super-resolved H2B-JF646 with diffraction-limited GFP-tagged BRD4. (d) Quantification of degree of clustering using the L-function (e) L-function for BRD4 mutants. (f) L-function after 8h exposure to 1uM (+)-JQ1 in DMSO and DMSO only. All scalebars 3um. ** p<0.01, *** p<0.001

### Heteropolymer model to simulate the interplay between condensates and chromatin

To interpret our experimental findings, we adopt a heteropolymer chromatin model^35^ to capture the interaction of chromatin with multivalent BRD4-like binders (Figure 4a). The heteropolymer consists of a coarse-grained bead-and-spring chain composed of *N*_*b*_ = 200 beads, connected by harmonic bonds with equilibrium length *r*_0_ whose energy is defined as

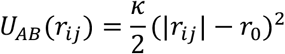

where *r*_*ij*_ is a vector connecting the center of a bead of type *i* to a bead of type *j* and *i, j ϵ* (*A, B*). In all simulations, we assume κ = 90*k*_*B*_*T*/*r*_0_^2^ where *k*_*B*_ is Boltzmann’s constant and *r*_0_ = 200nm. Random beads in the chain are selected to represent locally unacetylated (A-type particles) and acetylated chromatin (B-type particles). B-type particles undergo multivalent interactions with a third group of C-type particles, which can promote cross-linking of the polymer. We assume a Bernoulli probably of *p* = 0.3 for any given bead to be in an acetylated-like state. Interaction of multivalent chromatin binders with chromatin beads are then mediated by the following potential^36^:

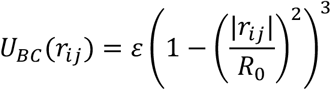

where *R*_0_ = 200*nm*. The potential *U*_*BC*_ is considered over a domain 0 ≤ |*r*_*ij*_| ≤ 2*R*_0_. In all simulations, ten replicates were run for each condition tested. A and B type particles within the chromatin polymer have repulsive interactions with *ε* = +10*k*_*B*_*T*. Binding energy of the acetylated beads with binders was varied with *ε*_*I*_ = 0*k*_*B*_ *T, ε*_*II*_ = −20*k*_*B*_ *T, ε*_*III*_ = −40*k*_*B*_*T* (Figure 4b). The dynamics of chromatin chains are approximated by Brownian dynamics within a cubic box with side length of 10um and periodic boundary conditions. Brownian dynamics follows the stochastic differential equation:

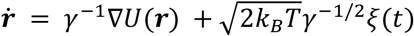

where γ is a diagonal friction tensor and *ξ*(*t*) is a three-dimensional delta-correlated white noise ⟨*ξ*(*t*) *ξ*(*t* + τ) ⟩ = δ(*t, t* + τ). We integrated the Brownian dynamics using *γ* = 10^−7^ and a time step of 10^−4^ sec over a 1 second duration. The first half of the simulation was discarded as burn-in. Integrating the Brownian dynamics showed an overall reduction in the diffusion coefficient *D* of single beads, and quasi-linear scaling of the diffusion coefficient with respect to temperature (Figure 4e).

**Figure 4:**
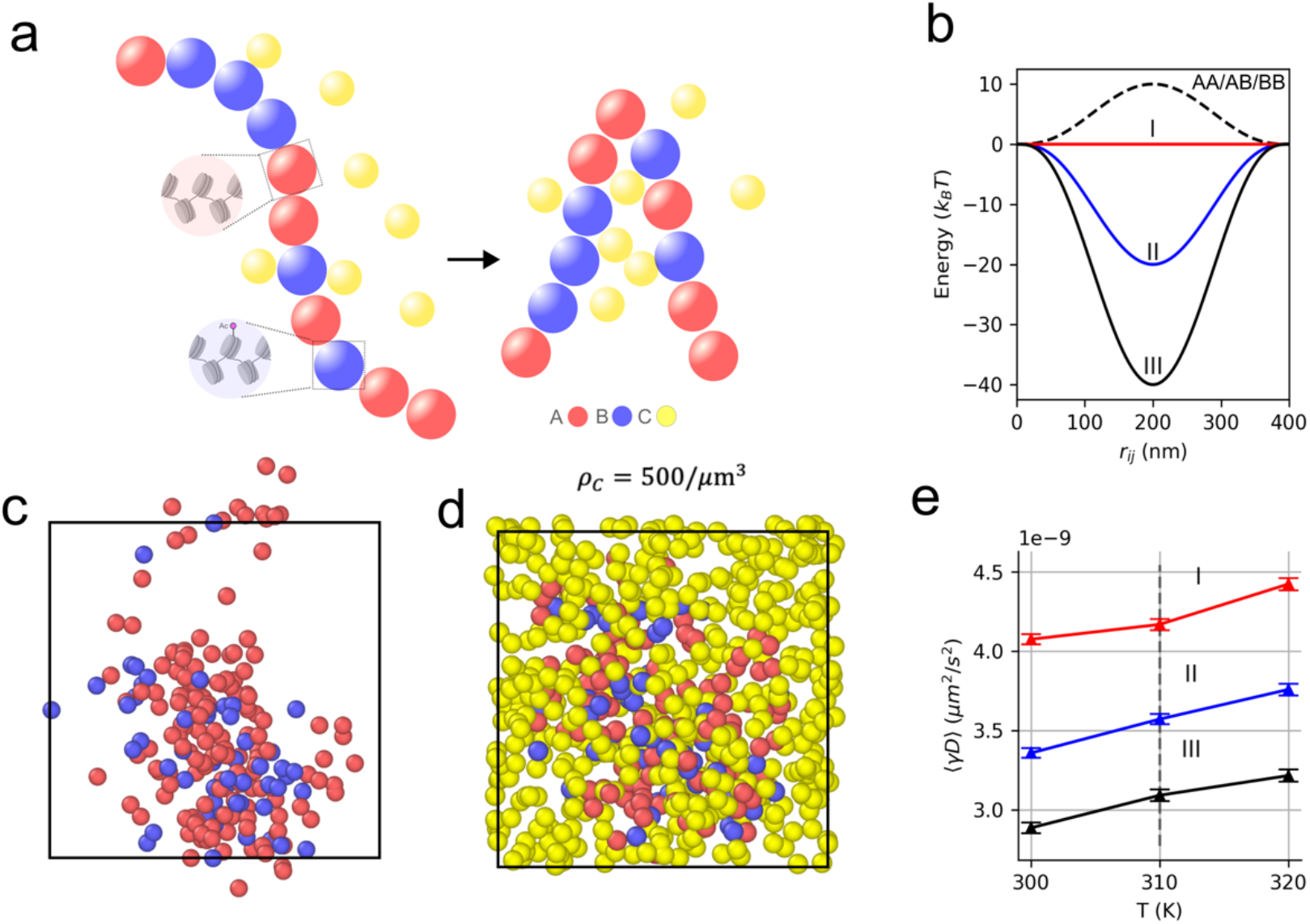
Multivalent chromatin binders reduce diffusion of nucleosome nanodomains. (a) Heteropolymer model of chromatin consisting of A-type, B-type, and C-type particles. (b) Interaction potentials *U*_*BC*_(*r*_*ij*_) of multivalent chromatin binders with B-type chromatin beads. (c,d) Example free heteropolymer and heteropolymer with a number density of C-type particles of ρ = 500/*μm*) in a 10um periodic box. (e) Scaled diffusion coefficient *D* for various chromatin binding energies of C-type particles, averaged over ten independent simulations, with burn-in discarded.

## DISCUSSION

Recent studies have demonstrated that BRD4 is present in discrete nuclear bodies, which exhibit properties of other well-studied biomolecular condensates, including rapid recovery of fluorescence after photobleaching and sensitivity to 1,6-hexanediol^25,37^. Both BRD4 long and short isoform are found in phase separated condensates in the nucleus and are associated with active gene transcription^3^. Importantly, CK2-mediated NPS phosphorylation regulates chromatin binding activity of BRD4^32^ as well as BRD4 phase separation^31^. This has led to the conclusion that phosphorylation of BRD4 inhibits interaction with chromatin and reduces phase separation, while remaining necessary for active gene transcription. Moreover, phosphorylated and unphosphorylated BRD4 form different molecular associations – transient polyvalent associations of unphosphorylated BRD4 contrast with the stable dimeric interaction and chromatin binding of phosphorylated BRD4^33^. Our data support the model that phosphorylated and unphosphorylated BRD4 form different molecular associations in the nucleus.

Reduced diffusivity of single nucleosomes for all mutants tested suggests a relationship between elevated colocalization and the degree of confinement of NN diffusion. At the same time, increased nucleosome diffusivity and reduced nucleosome clustering after (+)-JQ1 exposure indicate BRD4 condensate formation is necessary in maintaining the NN environment. Reduced diffusivity in the BD mutant is consistent with the colocazation result but indicate that the bromodomain mutations tested may alter chromatin and/or cofactor binding of BRD4 more subtle way that previously expected. We further hypothesize effects of BRD4 localization on the viscosity of the NN environment and potentially increased cross-linking of NNs. Indeed, we find this to be a natural result of molecular crowding resulting from overexpression of a constitutively phase separating protein, capable of multivalent interactions.

Substantial colocalization of the 7D mutant with NNs along with increased chromatin compaction also points towards a BRD4-mediated cross-linking mechanism of nucleosomes within the NN. This result of increased affinity can also be seen in coarse grained simulations of chromatin interactions with multivalent binders. We conclude that nascent BRD4 condensates are likely seeded by promiscuous interactions of BRD4 with cofactors, followed by phosphorylation and chromatin interactions mediated by the phosphorylated form. The stable dimeric interaction and binding of phosphorylated BRD4 to acetylated chromatin would then mediate control of the chromatin architecture by promoting cross-linking of the chromatin fiber and acting a molecular ‘bridge’ between transcriptional condensates with chromatin.

ACKNOWLEDGEMENTS

Authors would like to acknowledge the funding support from the National Institute of General Medical Sciences (NIH 1R35GM147412), National Institute of Diabetes and Digestive and Kidney Diseases (NIH 1R03DK135457) and the National Science Foundation (NSF 2239262).

## MATERIALS AND METHODS

### Cell lines, cell culture conditions, and transfection

HeLa cells were cultured in DMEM supplemented with 10 percent fetal bovine serum (Gibco) at 37C 5% CO2 in a humidified incubator. Cultures were tested routinely for mycoplasma contamination; all tests were negative. For super-resolution experiments, cells were seeded in a 35mm FluoroDish (WPI), and transfected using Lipofectamine 3000 with pBREBAC-H2B-Halo plasmid (Addgene plasmid #91564) (ThermoFisher #L3000008), pcDNA5-Flag-BRD4-7A (Addgene Plasmid #90006), pcDNA5-Flag-BRD4-7D (Addgene Plasmid #90007), pcDNA5-Flag-BRD4-BD (Addgene Plasmid #90005), pcDNA5-Flag-BRD4-WT (Addgene Plasmid #90331)

### Plasmid DNA Purification

Plasmids were transformed in E. coli at 4C and selected using an antibiotic agar plate. A single colony from the plate was selected and placed into sterile antibiotic LB Broth followed by incubation with shaking at 37C for 12h. After amplification, DNA was purified using a Miniprep kit (Promega). Following extraction, the concentration and purity were measured using the NanoDrop 2000 software. Plasmids were stored with optimal concentration and purity at -20C.

### Super-resolution imaging of nucleosome nanodomains in living cells

After transient transfection, H2B-Halotag HeLa cells were incubated with 3pM JF646 HaloTag ligand overnight. Cells were imaged in a dSTORM photoswitching buffer containing 100mM MEA, 50 ug/ml Glucose Oxidase, and 3.4 mg/ml Catalase (Sigma). Buffer pH was adjusted to ∼8 using HCl. Movies were collected using a custom Olympus IX83 microscope body equipped with an Olympus 60X 1.25NA oil-immersion objective. During imaging cells were maintained at 37C and 5% CO2 in a stage top incubator (Tokai Hit). Images were projected onto an ORCA-Fusion sCMOS camera (Hamamatsu) and 2000 frames were captured at 100fps. The microscope was controlled using Micromanager software. HaloTag-JF646 molecules were imaged using oblique illumination with a 640nm laser (Excelitas) held at 20mW, as measured at the back focal plane of the objective. Super resolution reconstructions were obtained using the ThunderSTORM ImageJ plugin. Background signal was subtracted using a rolling ball filter with radius of 10 pixels. Spots were fit using an integrated Gaussian point spread function model with maximum likelihood estimation^38,39^. Experimental conditions for single molecule tracking are nearly identical. However, H2B-Halotag HeLa cells were incubated with 3pM JF646 HaloTag ligand. HaloTag-JF646 molecules were illuminated at 10mW, 100 frames were captured at 10fps. All statistical analyses were performed with N=15-20 cells, per group.

After precise positions of the fluorophores are obtained, Besag’s L-function *L*(*r*) is used to analyze the clustering. The L-function is a transformation of Ripley’s K-function. The K-function is designed such that *K*(*r*) is the number of localizations within a radius *r* of a randomly chosen localization. Importantly, in the case of complete spatial randomness, *L*(*r*) = *r*. Thus, in general, to quantify the degree of clustering, one uses *L*(*r*) − *r*, which measures the deviation of a point pattern from complete spatial randomness.

### Colocalization of BRD4 mutants with nucleosome nanodomains

We colocalize FLAG-tagged BRD4 mutants with nucleosome NNs by simultaneous FLAG immunofluorescence with imaging of sparsely labeled of H2B-JF646. Puncta were detected in both channels using the Laplacian of Gaussian (LoG) detection algorithm to generate a multi-type point pattern. We then computed the nearest neighbor distance distribution as the distance from a random H2B-JF646 puncta to the nearest BRD4-FLAG puncta. Nearest distances values were pooled over N=10 cells for each mutant.

### Single molecule tracking

Nucleosomes were localized using an integrated Gaussian point spread function model with maximum likelihood estimation^38,39^ and tracked using TrackPy Python software. Trajectories lasting less than 80 frames were removed from further analysis. The individual mean squared displacement (MSD) is computed as follows.

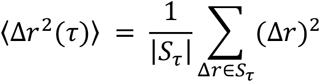

where *S*_*τ*_ is the set of all displacements in a time interval τ. The diffusion coefficient for both simulations as well as experimental data was computed by linear regression of the formula:

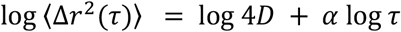

### Immunofluorescence

Cells grown in 35mm dishes were fixed with Formaldehyde in 1x PBS at 37C incubator for 20 minutes, and then permeabilized with 0.3 percent (v/v) Triton-X100 (Sigma-Aldrich) in PBS and blocked for 1h in 5 percent (w/v) nonfat dry milk at 4C. Cells were incubated overnight at 4C using primary antibodies anti-FLAG (Sigma-Aldrich, clone M2; 1:1000), and anti-BRD4 (Cell Signaling, clone E2A7X; 1:1000) in blocker. Secondary antibodies for BRD4 (Cell Signaling Anti-Mouse IgG-Alexa488, 1:1000) were used.

### Immunoblotting

Cells were washed and lysis buffer added (RIPA buffer: PMSF: protease inhibitor cocktail: orthovanadate=100:1:2:1). Cells were then scraped and sonicated for 15 seconds using an ultrasonic homogenizer. Lysate was centrifuged at high speed (13200r/min) for 15 minutes at 4C to pellet the cellular debris. Total protein concentration was determined by the BCA Protein Assay Kit (Thermo Fisher). For electrophoresis, protein samples were prepared according to a protein-4x loading buffer (containing DTT) ratio of 3:1, 4x loading buffer containing DTT was diluted with 3 aliquots of protein sample. The sample was mixed and vortexed, then heated at 95C for 5 min, followed by vortex and centrifuge. After running the gel, it was removed from the cassette and assembled inside the Trans-Blot Turbo Transfer System cassette. Transfer was run at 2.5A and 25V for 7mins. The sample was then blocked for at least 1 hour using 5% skim milk blocking solution prepared with PBS in RT. Primary FLAG antibody was diluted in PBST with 3% skim milk (1:500) and incubated at 4C overnight. The secondary antibody (Licor Anti-Mouse IgG-IRDye 800CW) was diluted in PBST with 3% skim milk (1:5000) and placed on a rocker and incubated in the dark at RT for 45min. Western blots on Nitrocellulose membranes were scanned using the Odyssey fluorescence scanning system software.

